# Kontextual: Reframing analysis of spatial omics data reveals consistent cell relationships across images

**DOI:** 10.1101/2024.09.03.611109

**Authors:** Farhan Ameen, Nick Robertson, David M. Lin, Shila Ghazanfar, Ellis Patrick

## Abstract

State-of-the-art spatial proteomic and transcriptomic technologies can deeply phenotype cells in their native tissue environment, providing a high throughput means to effectively quantify spatial relationships between diverse cell populations. However, the experimental design choice of which regions of a tissue will be imaged can greatly impact the interpretation of spatial quantifications. That is, spatial relationships identified in one region of interest may not be interpreted consistently across other regions. To address this challenge, we introduce Kontextual, a method which considers alternative frames of reference for contextualising spatial relationships. These contexts may represent landmarks, spatial domains, or groups of functionally similar cells which are consistent across regions. By modelling spatial relationships between cells relative to these contexts, Kontextual produces robust spatial quantifications that are not confounded by the region selected. We demonstrate in spatial proteomics and spatial transcriptomics datasets that modelling spatial relationships this way is biologically meaningful. We also demonstrate how this approach can be used in a classification setting to improve prediction of patient prognosis.

## Introduction

With the rapid uptake of spatial omic technologies such as MIBI-TOF, IMC, Xenium and MERFISH (Bressan et al. 2023), there exists an unprecedented opportunity to improve our understanding of how cells interact in their natural biological context. By coupling transcriptomic or proteomic profiles of cells with their location *in situ*, highly multiplexed spatial omics technologies enable the quantification of the spatial orientation of an unprecedented number of cell types simultaneously providing insights into complex cellular arrangements in a tissue (Zhang et al. 2024). These insights into cellular interactions can help reveal underlying biological processes, for example, in understanding the mechanisms of tumourigenesis and invasion (Lewis et al. 2021). For instance, tumours with high CD8+ T cell infiltration have been shown to be associated with positive survival outcomes in breast cancer (Mahmoud et al. 2011). More broadly, such interactions can be useful in determining spatial biomarkers that are able to predict patient prognosis and response to treatment (Wang et al. 2023, Nordmann et al. 2024). However, how best to analyse these complex high-dimensional data to facilitate robust interpretations of spatial patterns remains a challenge.

Currently spatial analysis methods can be predominantly grouped into two cate-gories, neighbourhood identification and spatial inference approaches. Neighbourhood identification approaches aim to resolve the complex tissue architectures imaged by these technologies into distinct cellular neighbourhoods, niches or spatial domains via novel dimension reduction approaches such as tensor decomposition, auto-encoders and clustering (Patrick et al. 2023, Schürch et al. 2020, Xu et al. 2022, Long et al. 2023, Shang & Zhou 2022). While these methods can identify intricate tissue structures, it is not always intuitive to assess how they are altered in the presence of disease. Alternatively, spatial inference approaches for identifying cell-to-cell interactions or cell localisation specifically test how two cell types are interacting with one another via Monte Carlo strategies or established spatial analysis techniques such as Besag’s L-function and Ripley’s K-function (Schapiro et al. 2017, Keren et al. 2018, Canete et al. 2022, Besag 1977). However an often overlooked aspect of current methods is they typically assume that cells are either distributed homogeneously across the field of view (FOV) of the image or that all cell types follow similar expected distribution under the null hypothesis of no interaction. This however is not always the case, some examples are paneth cells which are located at the base of the small intestines, podocytes which are found in the bowman’s capsule in the kidney, pneumocytes which line the alveolar compartment of the lungs (Lueschow & McElroy 2020, Reiser & Altintas 2016, Castranova et al. 1988). These cell types cannot exist outside of their local context, and thus is inappropriate to assume that these cell types can be found anywhere within the FOV of the image or exhibit similar distribution to the other cell types in the image. With the ability to appropriately define how cells are distributed in tissue under the null hypothesis of no interaction, the conclusions from spatial inference tools may be inappropriate or misleading, resulting in spurious significant associations. Thus there is a critical need for spatial analysis tools which allow you to explicitly define the expected behaviour of the cells when performing spatial inference.

With spatial inference tools which compute distances between cells, such as those that use Monte Carlo strategies, the K-function etc. the observed quantification of spatial relationships need to be interpreted relative to the spatial quantification of when they are distributed randomly and not interacting. More formally, under the null hypothesis of no interaction, the expected distribution of the cells exhibit complete spatial randomness (CSR), that is the cell types can appear anywhere within the context of the FOV. Therefore one means to control the null hypothesis and thus the expected distribution of the cells is by controlling the context in which the spatial relationship are evaluated. In this scenario, the null hypothesis states that the cell types exhibit CSR within the area defined by the context. Thus, evaluating spatial relationships under different contexts can enable novel spatial conclusions which cannot be ascertained when using the full FOV as a context (Figure 1a). A classic problem in geospatial statistics, known as the modifiable areal unit problem demonstrates how context selection can impact the results of statistical tests (Zormpas et al. 2023), however minimal evaluation of the impacts of this on spatial inference has been reported. Intuitively researchers know CSR is an unrealistic assumption, that cells won’t be spread throughout and image randomly, and that there are clear and expected structures in the tissue. With this in mind, great care is typically invested when designing experiments to either specifically select regions of interest (ROI) to image or opportunistically capture as much tissue structure as possible, without considering the implications that the selected FOV may have on spatial inference. One major consequence of this is that spatial relationships inferred in one patient image or FOV may not be comparable to other patient images, which may be testing entirely different null hypothesis. This highlights a critical barrier to identifying spatial biomarkers from cell localisation inference approaches which are consistent within and across patient cohorts.

**Figure 1:**
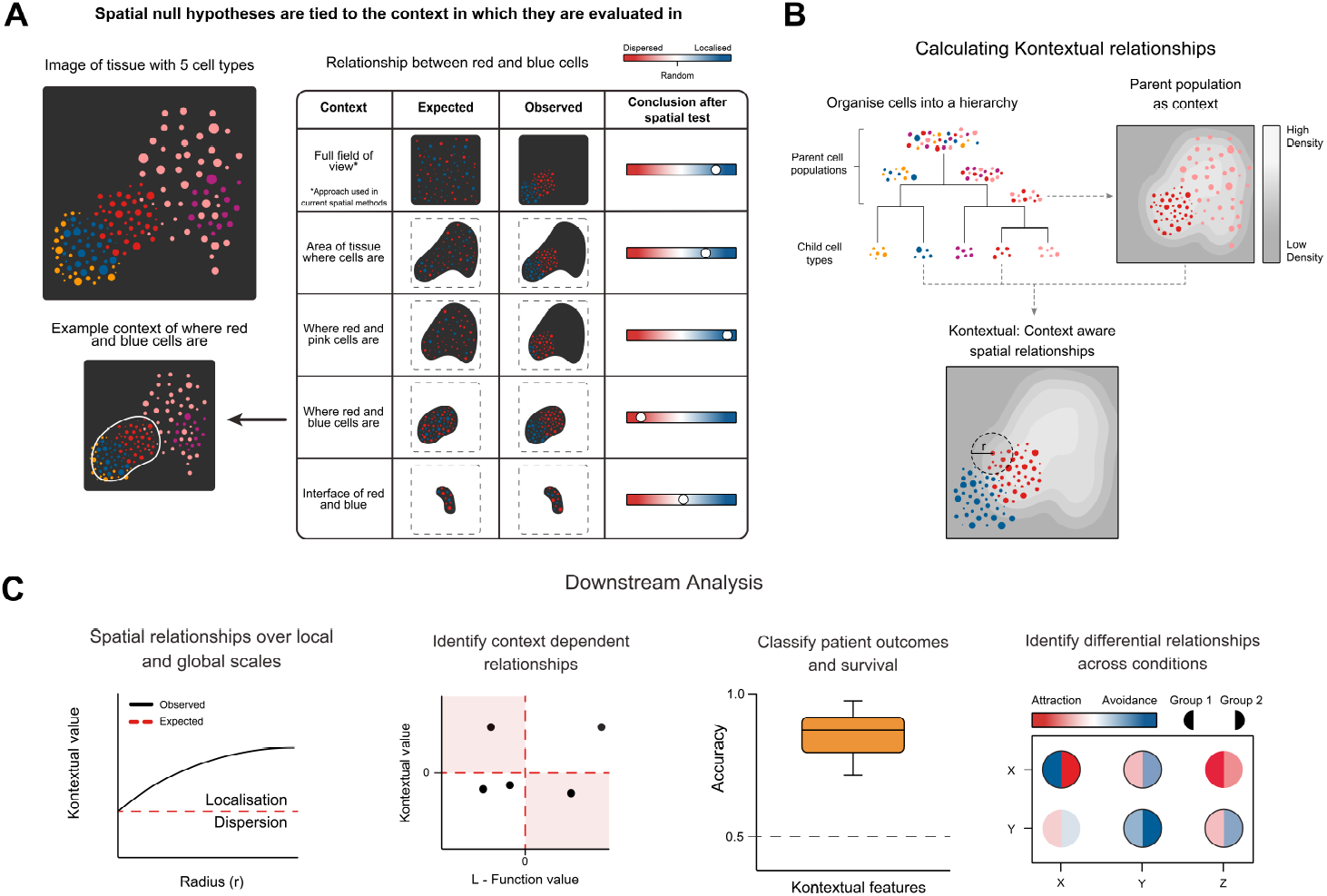
**(a)** The context in which spatial relationships are tested dictates the null hypothesis which is being tested. Thus testing the same spatial relationship between the red and blue cell types can lead to very different conclusions under different contexts. **(b)** A schematic overview of Kontextual. Kontextual models the spatial relationship between two cell types in the context of a parent population rather than the imaging area of the ROI. These parent populations are determined using a previously defined cell type hierarchy. By examining spatial behaviour relative to a parent, Kontextual provides robust spatial quantifications which are invariant to the choice of ROI and changes in tissue structure. **(c)** Kontextual features enable several downstream tasks. Such as evaluating cell type relationships on local and global scales, identifying relationships where incorporating contextual information enables novel conclusions, and identifying cell relationships associated with survival or patient outcomes.

To address this fundamental challenge, we have developed Kontextual, a novel analytical approach which evaluates cell type localisation within an appropriate context. Here we define a context as a region within each image which appropriately defines the expected distribution of the cells being evaluated, examples of contexts may include biological landmarks, spatial domains or groups of functionally similar cells. In this paper we exploit knowledge of cell function to produce these contexts, given the spatial distribution of functionally similar cells may model the expected distribution of a similar cell type than the entire FOV. By modelling spatial relationships between cells within biologically related context, we will demonstrate that Kontextual enables more appropriate spatial quantifications with clearly defined null hypotheses. Kontextual can be applied across multiple samples and thus used to 1) identify changes in spatial relationships between cell types that are associated with disease and 2) extract patient-level scores that can be further used in predictive or prognostic models. In this manuscript, using both spatial proteomic and spatial transcriptomic datasets we demonstrate Kontextual’s ability to identify novel cell type relationships with well defined null hypotheses, which cannot be captured by traditional spatial inference methods.

## Results

### Reframing the interpretation of cell-cell relationships with Kontextual

Here we present Kontextual, a method for performing inference on cell localisation which explicitly defines the contexts in which spatial relationships between cells can be identified and interpreted. Kontextual identifies pairwise cell type relationships that deviate from what is expected, using a broader population of cell types with similar behaviour as a reference. Briefly, Kontextual assumes that cells within an image have been segmented and are annotated with discrete cell type labels and x-y coordinates. It also requires either a user defined or data-derived cell hierarchy, grouping cells into functionally related populations. For various branches of the cell hierarchy, Kontextual calculates cell densities of the *parent* population of cells (Figure 1b). These densities are intended to produce a *context* which can be used to provide a reference for the expected spatial location of functionally related cells. Analysing cells as a Poisson point process, Kontextual then makes use of a modified inhomogeneous multitype K-function (Methods) to quantify cell type relationships within the context of the spatial behaviour of the parent population. We will demonstrate that this strategy produces robust quantification of cell type relationships which are invariant to complex tissue structures. Throughout the results section we demonstrate how the features generate by Kontextual affects the behaviour and interpretation of various downstream tasks (Figure 1c).

### Kontextual identifies cell relationships invariant to tissue structures in breast cancer images

Kontextual can quantify spatial relationships between cell types in tissue with distinct compartmentalisation. To illustrate this, we first selected a MIBI-TOF image acquired from a single triple negative breast cancer patient (Keren et al. 2018) that has a strong separation between immune cells and tumour cells (Figure 2a). By summarising each cell by an x-y coordinate of its centroid and a corresponding cell type label, the spatial relationship between P53^+^ tumour cells and immune cells can be visually assessed (Figure 2a). Qualitatively, we would argue that it is reasonable to make two conclusions. The first, that P53^+^ tumour cells are not mixing with immune cells, that is they are dispersed. The second, that P53^+^ tumour cells are closer to immune cells than other tumour cells. We will next show quantitatively that both of these conclusions are valid, but arise from different hypotheses.

**Figure 2:**
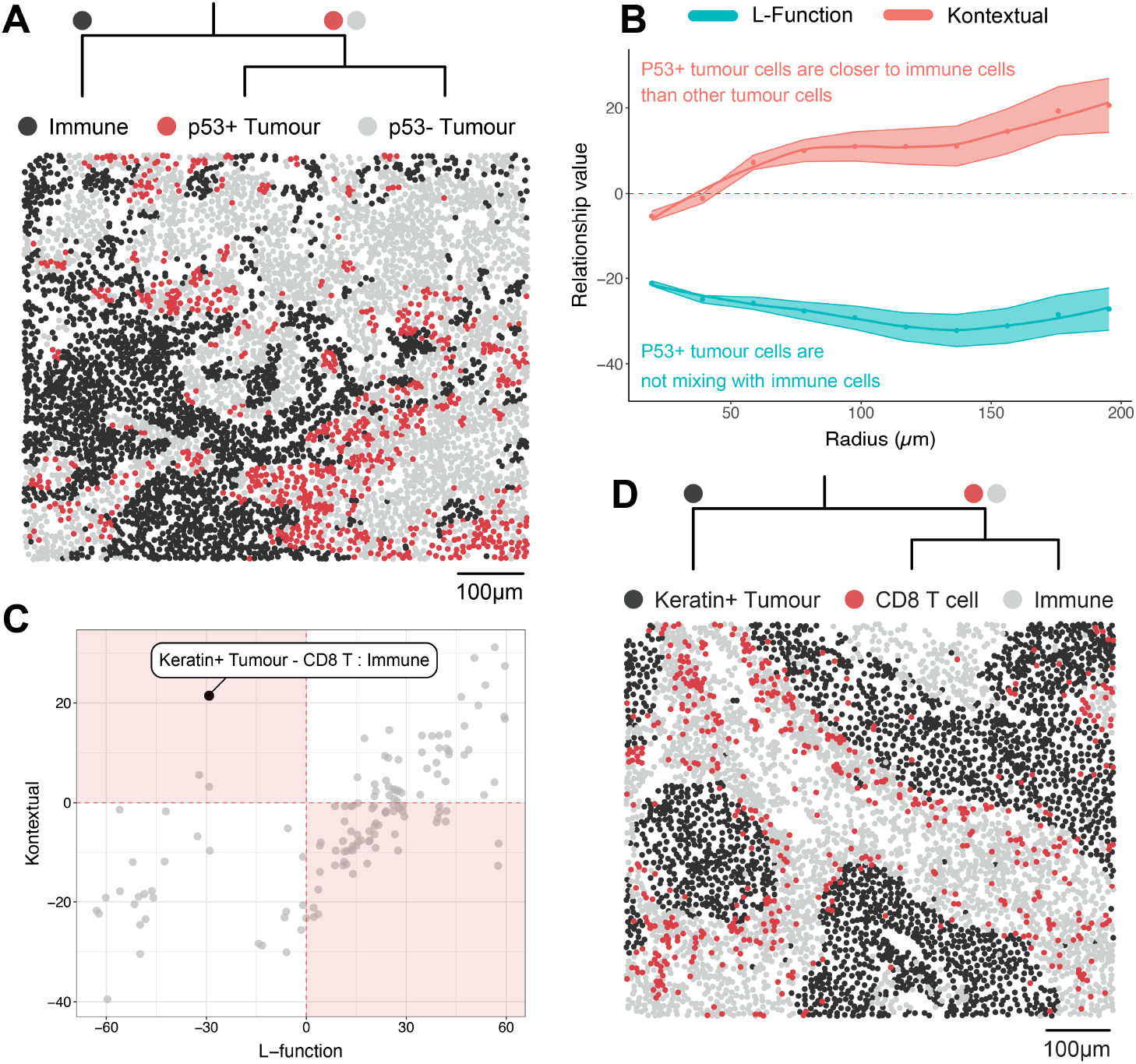
Kontextual models spatial relationships in the context of hierarchical cell structures. **(a)** Patient 6 image from Keren et al. (2018); P53+ tumour cells appear to be localised with immune cells after accounting for the context in which tumour cells reside in this image. (**b**) Assessing the relationship between immune cells and P53+ tumour cells in patient 6 image using the L-function and Kontextual. Positive points (above the red dashed line) indicate localisation, while negative points (below the red dashed line) indicate dispersion. (**c**) A scatter plot comparing the average L-function and Kontextual value of every pairwise cell type relationship across all breast cancer images. Points in the off diagonal quadrants (highlighted) represent relationships where accounting for a context results in differing conclusions. On such relationship is between Keratin^+^ tumour cells and CD8 T cells, the L-function describes this relationship as dispersed, however accounting for the context of immune cells Kontextual describes these cells as localised. (**d**) Example image highlighting the relationship between Keratin^+^ tumour cells and CD8 T from patient 5. CD8 T cells appear to not mix with Keratin^+^ tumour cells as confirmed by the L-function. However relative to the spatial behaviour of the immune cells, CD8 T cells appear to be localised and infiltrating the Keratin^+^ tumour body as identified by Kontextual.

To quantify the spatial relationship between P53^+^ tumour cells and immune cells we applied several established spatial inference techniques. These other techniques can be split into three broad categories: radius-based (Besag 1977, Baddeley et al. 2000, Canete et al. 2022), graph-based (Windhager et al. 2023) and density-based (Masotti et al. 2023). However, all are similar in the overall hypothesis tested; *‘acknowledging that two types of cells could be anywhere in the FOV of the image, are they closer or further away to each other than expected by chance?’*. The sign of the implemented radius-based methods are immediately interpretable with positive values indicating localisation between cell types and negative values indicating dispersion. The L-function (Besag 1977), inhomogeneous L-function (Baddeley et al. 2000) and spicyR (Canete et al. 2022) all quantify some measure of the average average number of cells of type A within a specified radius of cells of type B and produced values of −29.44, −21.93 and −26.24 respectively (Table S1) indicating dispersion. We applied a density based approach called DIMPLE (Masotti et al. 2023), where cell type densities are estimated using a kernel, and the distances between cell type densities are calculated to assess association. DIMPLE produced a value of −0.32 which indicates dispersion. We also implemented graph-based approaches using kNN, centroid expansion and Delaunay triangulation and (Windhager et al. 2023), resulting in observed values of 1.05, 0.02, 0.02 respectively. For these graph-based approaches, smaller values indicate dispersion and large values localisation. To ensure that we can obtain consistent interpretation of the test statistics for each of the spatial inference techniques, we performed permutation of cell type labels (n = 1000), and generated permutation-based Z-scores by subtracting the mean and dividing by standard deviation of the permuted test statistics. Upon examining these Z-scores, we observed that all Z-score values were negative (Table S1) indicating the observed relationship between P53^+^ tumour cells and immune cells are more dispersed than expected by chance. These results all support the valid conclusion that that P53^+^ tumour cells are not mixing with immune cells, which is consistent with the strong tissue compartmentalisation observed in the image (Figure 2a).

We next examined the behaviour of Kontextual, which can acknowledge the strong compartmentalisation of tumour cells observed in the image. Like the L-function and other radius-based approaches, Kontextual quantifies the spatial relationship between two cell types where positive (negative) values indicate that there are more (less) points of one type within radius *r* of another type. Using the parent population of ‘tumour cells’ as a context to perform spatial analysis, Kontextual produces positive values over a range of radii (Figure 2b). That is, acknowledging the context of where tumour cells are located in the image there is evidence of localisation between P53^+^ tumour cells and immune cells. For contrast, when applying the L-function to assess the relationship between P53^+^ tumour cells and immune cells over a large range of radii, we obtain negative values for all radii, providing evidence that these cell types are dispersed over both local and global ranges (Figure 2b). Summarising the results from these two different tests; when we expect that tumour cells could be anywhere in the image, P53^+^ tumour cells and immune cells are dispersed; when we provide the context of where we expect tumour cells to be in an image, P53^+^ tumour cells and immune cells are localised.

Evidently, discrepancies between the L-function and Kontextual can reveal cell type relationships which are context dependent. The conclusions made previously using the L-function and Kontextual are both valid, with the differences between these two methods attributed to the differences in the hypothesis which each method tests. By plotting the average L-function and Kontextual values of each cell type relationship calculated across all images, we can further examine where these metrics disagree (Figure 2c). To calculate Kontextual values a cell type hierarchy was created using prior knowledge of the biological relationships between the cell types (Figure S1). One such relationship, where the L-function and Kontextual disagrees, is between CD8 T cells and Keratin^+^ tumour cells. Ordinarily this relationship appears dispersed (L-function value = −29) as described by the L-function, however after accounting for the spatial context of where immune cells are located in the images, Kontextual provides evidence of localisation (Kontextual value = 21). This discordance in conclusion is illustrated in the patient 5 image (Figure 2d). Globally, the CD8 T cells are mixing less with the Keratin+ tumour cells than expected by chance, justifying the L-function’s negative value. However, in the context of where other immune cells are in the image, CD8 T cells are closer to, and infiltrating, the tumour border. By examining the differences between the L-function and Kontextual, we can identify cell type relationships, which otherwise would be confounded by the presence of strong tissue structures.

### Kontextual performs as expected in simulated data

To evaluate the behaviour of Kontextual in a controlled setting we applied Kontextual to simulated tissue images. Images containing a context dependent, context independent, and random cell type relationship were simulated using the spatstat R package (Baddeley & Turner 2005) (Methods). In the images containing a context dependent relationship, the L-function quantifies the relationship between *A* and *B* as dispersed, across all values of radii used, where *A* and *B* are two populations of cells. By accounting for the spatial behaviour of *A*’s context (*A* and a third population of cells, *C*) Kontextual identifies localisation between *A* and *B* (Figure 3). In images where a context independent relationship between *A* and *B* exists i.e. *A* is distributed similarly to the cells in its broader context, both the L-function and Kontextual quantify dispersion between *A* and *B*. Finally, in images where all cell types are completely randomly distributed in the image, both the L-function and Kontextual return values close to 0, indicating complete spatial randomness. Therefore, using simulated images, we have illustrated the expected behaviour of Kontextual when context dependent relationships do and do not exist.

**Figure 3:**
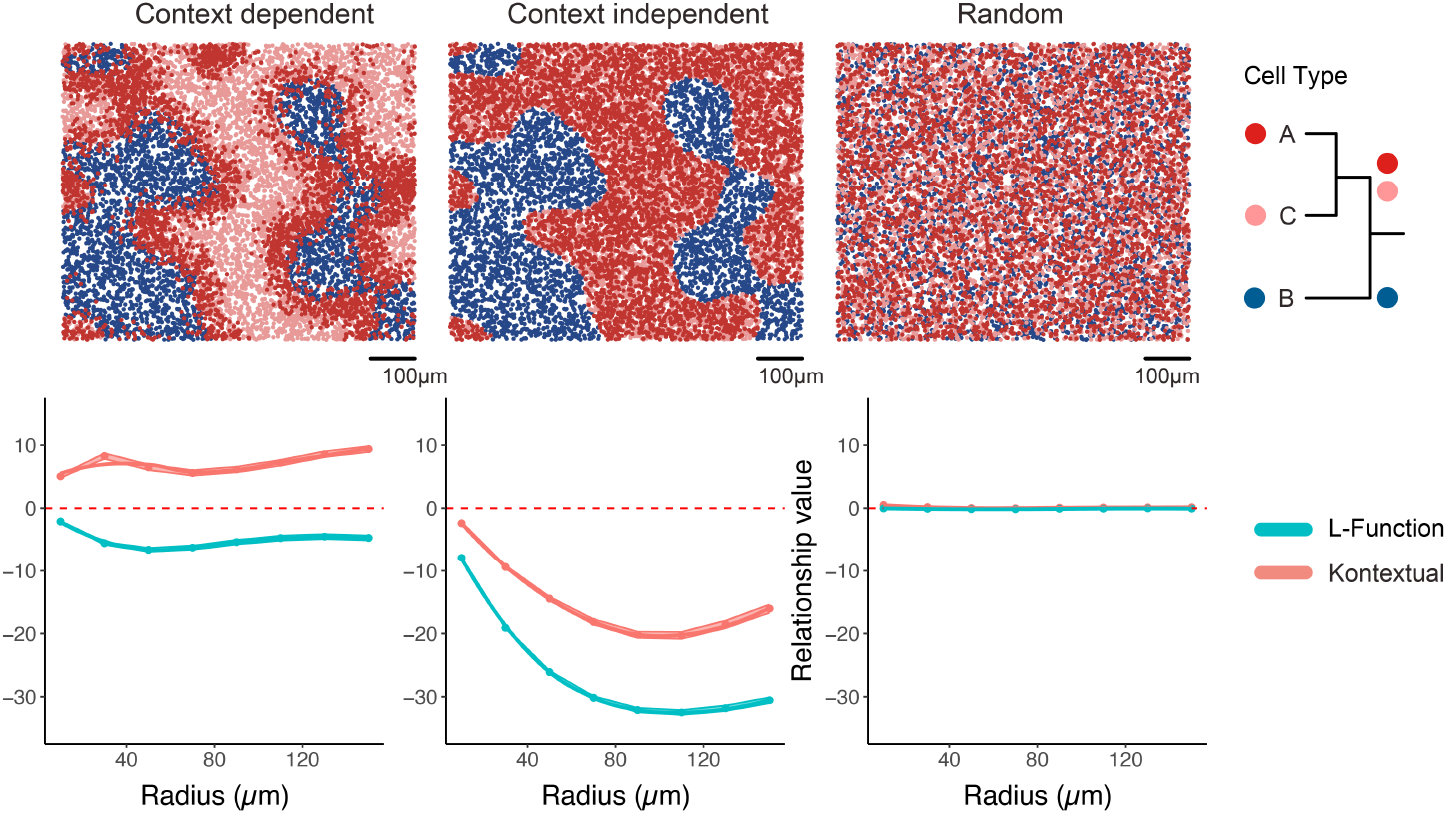
**Top row:** Examples of simulated images containing a context dependent, context independent and random relationship between cell type A and B. **Bottom row**: Assessing the relationship between the cell type A and cell type B over a range of radii using the L-function and Kontextual. Points above the red line indicate localisation, points below the line indicated dispersion. In the images where the spatial relationship between A and C are context dependent the L-function and Kontextual disagree, when this relationship is context independent there is agreement between Kontextual and the L-function. When the cell types are randomly distributed in the image, both the L-function and Kontextual produce values close to 0 indicating no relationship.

### Comparison of Kontextual with Monte Carlo approach

Next we evaluated the computational performance and the interpretation differences between Kontextual and a Monte Carlo based approach employed in Keren et al Keren et al. (2018). The Monte Carlo approach computes the average distance between *A* and their nearest *B*, cell labels are repeatedly randomised within *A*’s context and distances between *A* and *B* are recalculated to form a null distribution. By comparing the original distance between *A* and *B* to the null distribution, this approach can quantify how the behaviour of *A* deviates from the cells in its context. When applied to a single simulated image of size 800µm^2^ containing 12000 cells, the Monte Carlo approach takes 28.26 seconds using 1000 permutations, Kontextual using a radius of 50µm performs 10 times faster taking 3.11 seconds. Moreover, there are nuanced differences in the insights derived from using Kontextual versus the Monte Carlo approach. A key distinctions lies in how each method considers the spatial context: Kontextual acknowledges the spatial behaviour of the context, whereas the Monte Carlo approach conditions on the context. Specifically, the Monte Carlo approach assesses whether cell type *A* aligns or diverges from the spatial distribution of its context. In the context dependent image, the Monte Carlo approach states on average cell type *A* is 16.11µm closer to their nearest cell type *B* compared to its context (Figure S2), this is concordant with the findings from Kontextual. However, in the images with context independent (distance = −0.15µm) and random relationships (distance = −0.03µm), the Monte Carlo approach shows that cell type *A* exhibits the same spatial behaviour as its context, which does not elucidate the specific relationship between *A* and *B*. By acknowledging the context rather than conditioning on it, Kontextual is able to ascertain the relationship between *A* and *B* within the context that *A* belongs to.

### Kontextual features predict survival in breast cancer

To determine if spatial features generated using Kontextual capture meaningful biological signal, we performed a survival analysis on 33 triple negative breast cancer patients (Keren et al. 2018) (Methods). We first examined if the Keratin^+^ tumour and CD8 T cell relationship found earlier was associated with survival using a Cox proportional hazards model. Using immune cells as a context, the localisation between CD8 T cells and Keratin+ tumour cells is associated with longer survival (P-Value = 0.035) (Figure 4a). Alternatively, without using the expected locations of immune cells as a context for reference, the spatial relationship between Keratin^+^ tumour cells and CD8 T cells quantified by the L-function are not associated with differences in survival (P-Value = 0.82).

**Figure 4:**
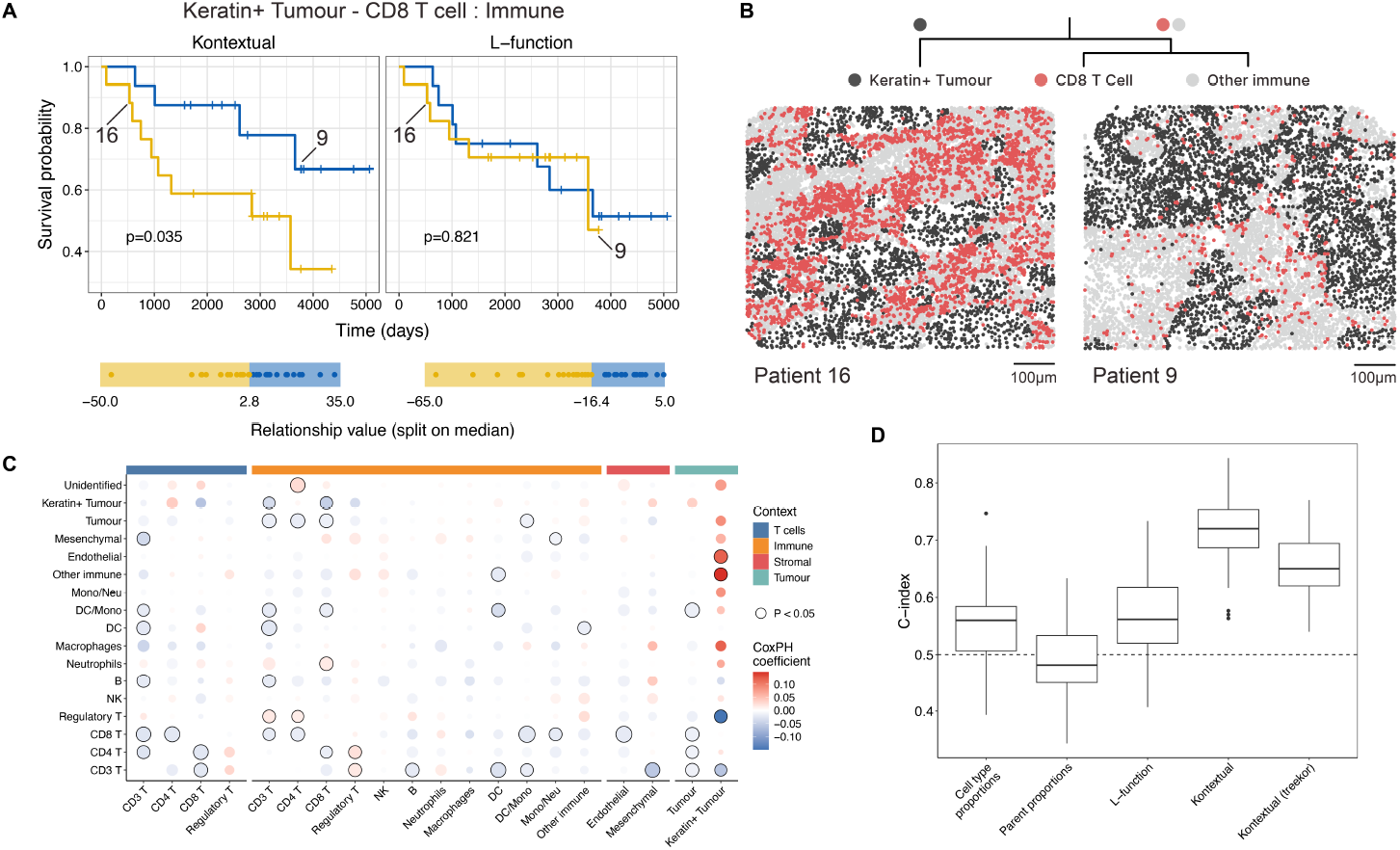
Survival analysis and classification using Kontextual. (**a**) Kaplan Meier curves of the relationship between Keratin+ tumour and CD8 T cells with immune as the context calculated using Kontextual and the L-function across all patients. The curves are stratified by the median value of each method. The reported p-values are from a Cox proportional hazard model fitted on the original feature values prior to splitting them on the median. (**b**) Two compartmentalised tumour images from patient 16 and 9. The L-function cannot is not able to distinguish between these images labelling the relationship between CD8 T cells and Keratin^+^ tumour cells as dispersed. By using immune cells as a context, Kontextual identifies CD8 T cell infiltration into the tumour body in patient 9 but not patient 16, and is therefore able to stratify survival outcomes much better than the L-function. (**c**) Bubble plot depicting the Cox proportional hazard coefficient for every pairwise Kontextual feature. Outlined circles represent P < 0.05, a positive coefficient suggests increased localisation between the cell types is associated with poorer survival outcomes, and vice versa. (**d**) Survival classification results, using regularised Cox Regression models trained on spatial and non spatial features. The models were evaluated using cross-validation and performances assessed using Harrell’s C-index, where a higher C-index indicates better survival classification performance.

According to Keren et al., patients with compartmentalised tumours exhibit improved survival than those with mixed tumours. Our results suggest that survival within compartmentalised tumours can be further stratified based on CD8 T cell infiltration into Keratin^+^ tumours. For example, while patient 16 and 9 both have strongly compartmentalised tumours (Figure 4b), they have very different survival times: 530 vs 3,767 days respectively. Kontextual quantifies the CD8 T cell and Keratin^+^ tumour relationship as dispersed in patient 16 (value = −12) and localised in patient 9 (value = 18), suggesting that increased CD8 T cell infiltration into Keratin^+^ tumours is associated with positive survival outcomes. The increased CD8 T cell infiltration in patient 9 can be visually confirmed in the images (Figure 4b) The strong compartmentalisation in these images make it difficult for the L-function to quantify this CD8 T cell infiltration, labelling both the patient 16 (value = −37) and 9 (value = −19) images as dispersed (Figure 4a). In this example the L-function correctly identifies that both images represent strong compartmentalised tumours. However beyond this conclusion, the L-function is not able to elucidate the subtle relationships between CD8 T cells and Keratin^+^ tumours without further contextual information. By leveraging Kontextual to consider the immune context, we uncover a pertinent spatial relationship related to survival that would have been missed if performing a context agnostic spatial analysis.

While we have identified a single spatial relationship associated with survival, we next looked to see if we could use all of the relationships in the datasets to predict patient survival. Testing every pairwise combination of cell types with four different contexts, Kontextual identified several features that were associated with patient survival (P-Value < 0.05) (Figure 4c). Next, we used a regularised Cox Regression model to predict survival using Kontextual values as predictors. This proved to be effective, producing an average cross-validated C-Index of 0.72, where higher values indicate better ability to predict survival. For this particular dataset, Kontextual was able to identify predictive signal that was not evident for the L-function (C-Index = 0.57) or cell type proportions (C-Index = 0.55) (Figure 4d). Further, when the contexts are determined in a data-driven way using a hierarchical clustering approach as calculated by the treekoR R package (Chan et al. 2021), survival prediction is also better than cell type proportions and the L-function. Overall we find that in this dataset, spatial features which incorporate hierarchical context information has superior predictive signal compared to classical spatial and non spatial quantifications.

### Kontextual identifies conserved relationships in large scale images

Finally we evaluated if the Kontextual features are applicable to a large scale imaging data sample. To this end, we applied Kontextual to a Xenium breast cancer image (Figure 5a) from Fu et al., which is 64 times larger (7500 x 5500µm) than the MIBI-TOF image (800µm^2^). When applied to this Xenium image, Kontextual also identified the spatial relationship between tumour cells and CD8 T cells (Figure 5b). We observed that CD8 T cells appear to aggregate and infiltrate the tumour body in this image (Figure 5c). Further, the additional RNAs quantified by Xenium allowed the tumour cells to be stratified into three subtypes; DCIS 1, DCIS 2 and invasive tumour. A comparison of the CD8 T cell interaction with the three tumour subtypes within this image revealed discrepancies between the quantifications using the L-function and Kontextual with immune cells as context (Figure 5d). In this case we observe the Kontextual metrics are more consistent with what we see visually in the image, where CD8 T cells aggregate around each tumour sub type (Figure 5e). Through its application on a Xenium image we show Kontextual is able to identify relationships in large scale images that are concordant with images of smaller scale.

**Figure 5:**
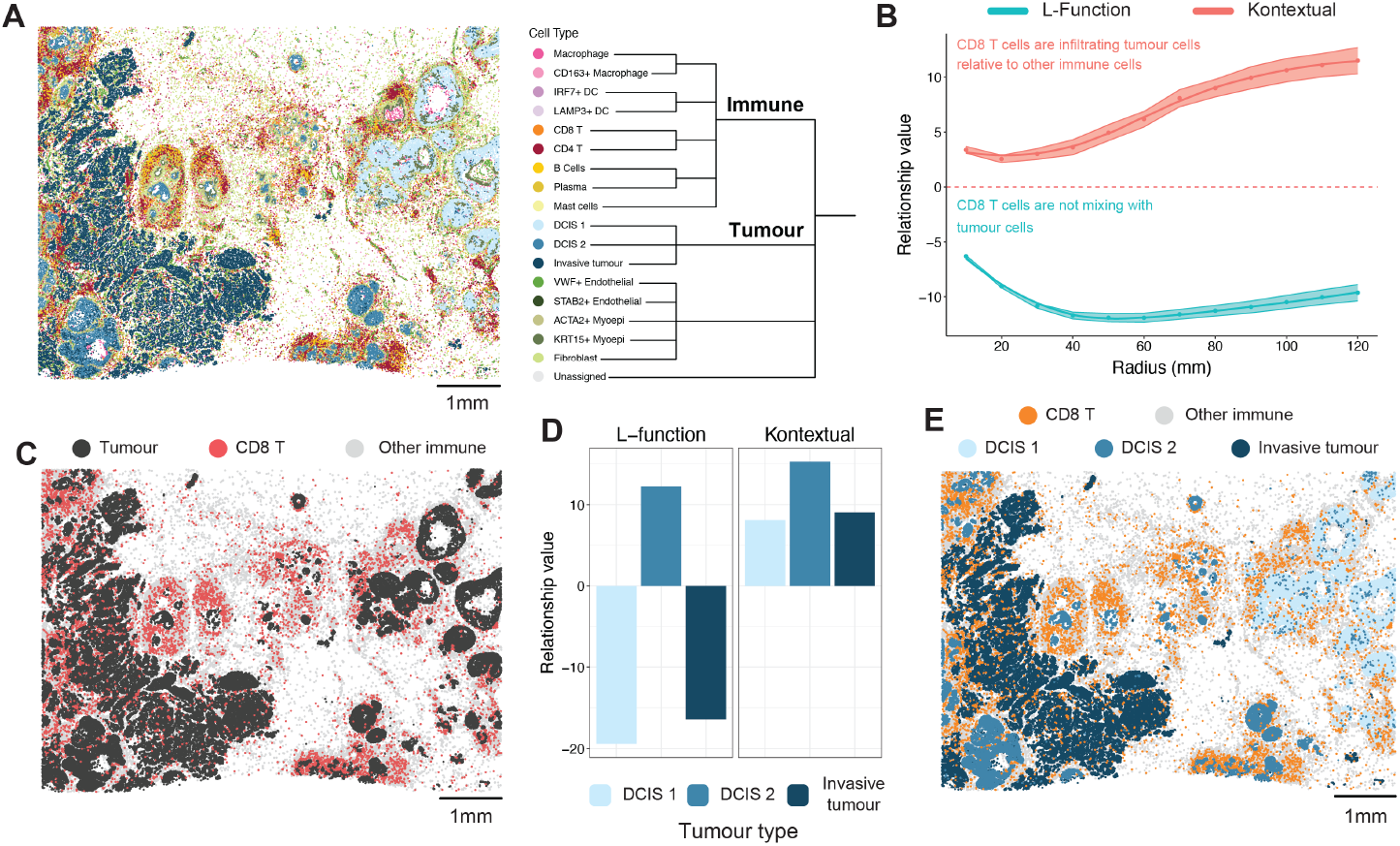
Kontextual applied to a Xenium breast cancer image. (**a**) Xenium image of breast cancer. The cell types are organised into a hierarchy based on biological function. (**b**) Examining the relationship between CD8 T cells and tumour cells over a range of radii using the L-function and Kontextual. Points above the red line indicate localisation, points below indicated dispersion. Kontextual indicates CD8 T cells are localising with tumour cells in the context of all immune cells. (**c**) Xenium image highlighting the relationship between CD8 T cells and tumour in the context of immune cells. We see CD8 T cells conglomerate around the tumour body much more than other the immune cells. (**d**) Comparing the L-function and Kontextual values examining the relationship between CD8 T cells and each sub-type of tumour. Kontextual identifies localisation (positive values) between the CD8 T cells and each of the tumour subtypes. (**e**) Xenium image highlighting the relationship between CD8 T cells and each tumour sub-type in the context of immune cells.

## Discussion

Here we have presented Kontextual, a novel analytical approach to quantify the localisation between two cell types accounting for the context in which the cell types appear. Kontextual produces quantifications of spatial relationships between cell types that are not confounded by the presence of distinct tissue structure. By incorporating the information on functional similarities present in hierarchical cell trees, Kontextual is able to quantify subtle cell relationships, such as the tumour infiltrating CD8 T cells, which current spatial inferences tools such as Monte Carlo, the inhomogeneous L-function, spicyR, DIMPLE, and several graph based approaches fail to detect. In this paper, we have illustrated the behaviour of Kontextual through its application in selected triple negative breast cancer images, and explored Kontextual’s statistical properties using a simulation study. Our findings demonstrate that Kontextual features are able to capture spatial relationships associated with patient survival, as demonstrated through survival analysis and classification performance. Finally, we demonstrate Kontextual is suitable for large scale images through its application to a Xenium breast cancer image.

Overall, it is important to note that all spatial inference methods that we applied generate valid results on the spatial relationships being evaluated, however Kontextual is distinct in terms of the null hypothesis which it tests. For example, in the Keren et al Patient 6 image (Figure 2a) the established spatial methods provide evidence that P53^+^ tumour cells are dispersed from immune cells, however, Kontextual provides evidence that P53^+^ are localising with immune cells relative to the tumour context, both interpretations are visually supported. The differences between existing methods and Kontextual are attributed to the subtle differences in hypothesis which each approach tests. Current spatial inference methods take a context agnostic approach, which assumes under the null hypothesis that any cell can appear anywhere in FOV of the image, in other words the null distribution of every cell exhibits CSR. However this assumption may not always align with a hypothesis a researcher wants to assess. For example, it is reasonable that a scientist would want to acknowledge that P53^+^ tumour cells are more likely to exist within the tumour region of the tissue and not in the immune region, which is counter to the assumption of CSR. Therefore compared to the null where P53^+^ tumour cells exhibit CSR, the existing spatial methods observe dispersion between P53^+^ tumour cells and immune cells as a result of the strong tissue structure. Alternatively, Kontextual assumes under the null hypothesis that the child cell type exhibits CSR only in the regions defined by its context. In this case, when the contexts represent a parent population in the cell hierarchy, Kontextual examines how the child cell deviates or aligns with its expected distribution as characterised by the spatial behaviour of its parent. Given the formulation of Kontextual, the L-function is a special case of Kontextual which uses the full FOV of the image as the context.

The choice of context is very important when using Kontextual. The context defines the expected distribution of the cell types under the null hypothesis. In our approach we select a context informed by canonical hierarchical groupings of cell function, that is, determined by prior knowledge. This is motivated under the idea that a parent population of functionally similar cells provides and appropriate and interpretable context to model the expected distribution of its child cell types, this assumption is also made in traditional suspension cytometry analysis (Chan et al. 2021). This assumption is valid in certain scenarios, such as in the Patient 6 breast cancer image (Figure 2a), where P53^+^ tumour cells are less likely to exist outside the space occupied by the tumour parent context. However these contexts may be inappropriate when the parent populations are sparse, leading to unstable estimations of the parent densities. Further work could examine the influence of the context selection according to differing sources of prior information in terms of consistency of the Kontextual quantifications and conclusions being drawn. It is thus important for users to choose a context which appropriately models the expected distribution of the cell types being evaluated, based on their biological knowledge and question.

It is worth noting that users are not limited to using hierarchical cell structure to define contexts. Contexts may be chosen based on biological factors such as presence of a certain biomarker, specific landmarks in the image, or selected regions of interest (ROI) within the image. Alternatively, users may opt to use a data-driven approach for defining contexts, as we did using the treekoR package (Chan et al. 2021). When used for predicting survival within in the MIBI-TOF breast cancer cohort, this data-driven approach for defining contexts using treekoR nearly performed as well as using prior knowledge. Further to this, the Kontextual approach could be further extended by using hierarchical pruning strategies (Huang et al. 2021), which could provide the most informative cell relationships and parent contexts within a particular cell hierarchy. Ultimately, regardless of approach, the chosen context has implications on the inferences and interpretations that can be made.

Another key consideration for Kontextual is the choice of radius to evaluate the cell relationships over. A small radius examines relationships over local distances and a larger radius examines relationships over a global scale. However, larger radii can introduce edge effects when the radius extends beyond the border of the image. Although Kontextual employs border correction to mitigate this, an excessively large radius will still lead to inaccurate quantifications of the spatial relationships due to extremely pronounced edge effects. Thus users should choose a radius appropriate to their biological questions. As demonstrated by our figures, we advocate that users examine spatial relationships over a range of radii when interpreting spatial relationships.

In this paper, we have presented Kontextual, a method to evaluate cell localisation relative to explicitly defined contexts. Kontextual can be used to compliment other spatial and non-spatial analysis techniques, producing feature sets which account for tissue structures present in images. It is our experience that all quantification approaches in a typical spatial analysis pipeline provide value. Here we have demonstrated that Kontextual can showcase unique relationships in a dataset that are necessary to paint a comprehensive biological picture with spatial omics data.

## Data and code availability

All datasets used in this study are publicly available. The MIBI-TOF triple negative dataset was obtained from the SpatialDatasets bioconductor package (Ameen et al. 2023). The Xenium dataset and all the code used to generate the results and figures are available on Zenodo: https://zenodo.org/doi/10.5281/zenodo.13308969. Kontextual is available in the Statial R package found on bioconductor: https://bioconductor.org/packages/release/bioc/html/Statial.html

## Acknowledgements

The authors thank all their colleagues, particularly at the Sydney Precision Data Science Center for their support and intellectual engagement. Special thanks to Sourish Iyengar and Alex Qin for their contributions to the Statial package. Thanks also goes to Dinny Graham for providing insights into the tumour sub types within the Xenium breast cancer image.

## Methods

### Overview of Kontextual

Kontextual quantifies the degree to which two populations of cells are localised in an image relative to a reference population of cells (Figure 1b). As input, Kontextual expects spatial profiling data that has been segmented and cell types labelled such that cells can be summarised by the x-y coordinates of their centroid and discrete cell types labels. It also assumes that a set of cell type names can be used to define a context population. Kontextual significantly extends on the multitype K-function, which we describe in further detail below.

### The multitype K-function

Cells can be modelled using a Poisson point process if their spatial location is simplified to a single x-y coordinate. Ripley’s K-function (Ripley 1976) is a well established tool for quantifying the spatial localisation of a Poisson point process. The multitype K-function can be used to quantify the spatial association between any two pairs of cell types *a* and *b* (Ripley 1976) where

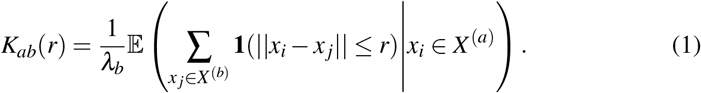

*K*_*ab*_(*r*) is the expected number of cells of type *b* lying within a distance *r* of a typical cell of type *a*, scaled by the intensity of cell type *b*. Where *X* ^(*a*)^ and *X* ^(*b*)^ represent all the cells of cell type *a* and *b* respectively, *x*_*i*_ and *x* _*j*_ represent the *i*^*th*^ and *j*^*th*^ cell in *X* ^(*a*)^ and *X* ^(*b*)^ respectively. The distance between every cell type *a* and *b* is represented by ||*x*_*i*_ − *x*_*j*_||, where if the distance between the two cell types falls below a radius *r* then 1 is added to the overall equation. This distance calculation is evaluated over every cell of type *a* and the total is divided by the total number of *a* cells. This produces the average number of cell type *b* with radius *r* of cell type *a*. Finally this value is divided by *λ*_*b*_, where *λ*_*b*_ is the global intensity of cell type *b*, such that

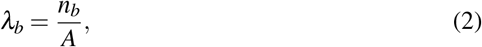

*n*_*b*_ is the total number of cells of type *b*, and *A* is the area of the image. Dividing this average by *λ*_*b*_ allows us to compare point patterns with differing intensities as the expectation of *K*_*ab*_(*r*) is *πr*^2^ if there is no spatial association between the cell types.

Following this definition of the multitype K-function, its observed value can then be calculated as

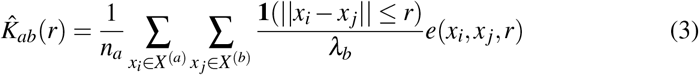

where *e*(*x*_*i*_, *x* _*j*_, *r*) is an edge correction weight. To simplify its interpretation, the multitype K-function will often be transformed into the centred Besag’s L-function (Besag 1977):

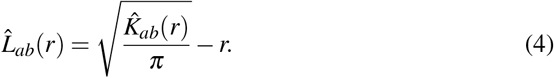

When there is no spatial association Besag’s L-function will be centred on zero and have approximately constant variance over *r*.

The multitype K-function can be extended to account for inherent tissue structure. The inhomogenous multitype K-function can be used to identify spatial associations that are independent of tissue structure and can be calculated as

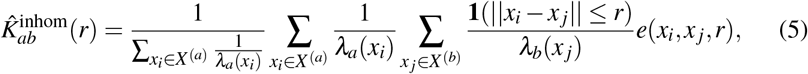

where *λ*_*a*_(*x*_*i*_) is the local intensity of cell type *a* at point *x*_*i*_, and *λ*_*b*_(*x* _*j*_) is the local intensity of cell type *b* at point *x* _*j*_. The local intensity of any cell type *c* at point *x*_*k*_ is defined as

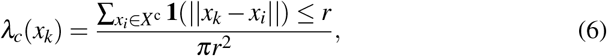

where the local intensity of a cell type *c* at point *x*_*k*_ represents the count of all cell type *c* with a radius *r* of *x*_*k*_, divided by the area of the circle *πr*^2^. In the inhomogeneous multitype K-function (Equation (5)) each point belonging to cell type *a* and *b* are therefore down-weighted by their local intensities. Doing so ensures that areas with high numbers of cells due to tissue inhomogeneity do not artificially inflate the K function value.

### Constructing Kontextual

Here we propose *Kontextual*, an extension of the multitype K-function as

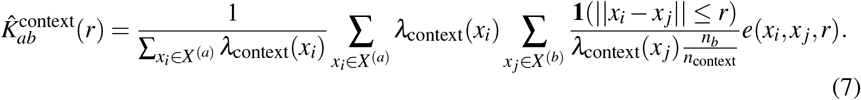

We introduce *λ*_context_, where context refers to the intensity of all cells belong to a particular context rather than just the individual intensities of a cell type. These contexts may be defined by pathologists as a region of interest or based on biological structures such as a tumour, or they may be defined via a data driven approach.

To account for the behaviour of cells relative to the context, the weights of cell type *a* and *b* are sampled from the context intensity (*λ*_context_), calculated using (Equation (6)) with all cells belonging to the context. Weighting the average over cell type *a* by the intensity of the context is used to restrict the average to cells within the context of interest. Similarly, dividing by 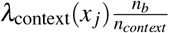 is used to scale the sum by the expected intensity of cell type *b* within the context. Note, cell type *b* must belong to the context, otherwise down-weighting the equation by *λ*_context_(*x* _*j*_) ill create a 0 division error. In addition to this, although cell type *a* may not explicitly belong to the context, *λ*_context_(*x*_*i*_) can still be calculated as it examines cells of type *a* which are found within *r* of a cell belonging to the context, as defined in (Equation (6)). Because cell type *b* belongs to the context, and Kontextual counts the number of cell type *a* within *r* of cell type *b*, the context intensity of cell type *a* (*λ*_context_(*x*_*i*_)) will always defined, if there exists no cell type *a* within *r* of cell type *b* then Kontextual will return NA. Finally the 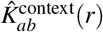 is transformed using Besag’s L transformation (4) to obtain the final Kontextual value.

### Defining tissue contexts using cell function relationships

In this paper we propose defining contexts that are analogous to the contexts that are used in suspension cytometry. In the analysis of suspension cytometry technologies such as CyTOF and flow cytometry, cell types are often defined via manual gating strategies or trees. In these gating strategies a *parent population* of cells is segregated into smaller child populations. When tasked with identifying if the proportion of cells in the child populations are changing between samples, these proportions are typically calculated relative to the parent population as opposed to the total population of cells. This arguably can improve robustness and interpretation of changes (Chan et al. 2021). As such *λ*_context_ is defined by the cell types belonging to a parent population using (Equation (6)), where cell type *b* is a child cell type of that parent.

Two strategies were employed to create the cell hierarchies required for Kontextual in this manuscript. The first strategy utilised biological understanding of the cell types to form hierarchies (Figure S1a), e.g. all tumour cells were grouped under one tumour parent population etc. The second strategy used a HOPACH hierarchical clustering strategy to create parent populations (Figure S1b), this was done using the treekoR R package (Chan et al. 2021).

### Application to breast cancer images

Kontextual was applied to a study aimed to characterise the tumour-immune microenvironment in triple negative breast cancer (TNBC) Keren et al. (2018). The dataset contains 38 TNBC images profiled using multiplexed ion beam imaging by time of flight (MIBI-TOF). A 36 antibody panel was used to capture multiple immune and tumour cell populations. Data of 197k segmented cells was downloaded from Keren et al.. In addition to the images, clinical information such as patient tumour grade, survival days and tumour type were provided. Analysis of patient tumour type constituted a majority of their analysis and the tumours were split into 3 classifications: cold, mixed and compartmentalised tumours. In addition to this, we applied Kontextual to a single multiplexed breast cancer image obtained using the 10x Genomics Xenium platform. Data was downloaded from (Fu et al. 2024) profiling 103K cells with cell type annotations corresponding to 18 distinct cell types. Hierarchical cell context was taken from the scClassify results (Lin et al. 2020) using a scRNA-seq dataset as reference.

### Comparisons to other methods

Konextual was compared to several other methods. These methods include the L-function (Besag 1977), inhomogeneous L-function (Baddeley et al. 2000), spicyR (Canete et al. 2022), graph approaches such as kNN, centroid expansion and Delaunay triangulation (Windhager et al. 2023), and DIMPLE (Masotti et al. 2023). The inhomogeneous L-function was calculated using a radius of 100*µm*, spicyR was calculated using multiple radii of 25, 50, 75, 100*µm*. The kNN graph was constructed using *k* = 20, centroid expansion used a threshold of 20, and Delaunay triangulation used a maximum distance of 20. DIMPLE used the default pixel size of 10 and a bandwidth size of 30 to calculate the kernel density and the relationship between the kernels was calculated using correlation. All methods were used to assess the spatial relationship between P53^+^ tumour cells and immune cells in the Keren et al. patient 6 image. To calculate the mean and standard deviation of each metric the cell type labels were permuted 100 times and each metric was recalculated. The Z-score was calculated by subtracting the permuted mean from the observed value and dividing by the permuted standard deviation.

### Simulations to assess performance of Kontextual

All simulated images were generated using the spatstat R package (Baddeley & Turner 2005). For each image, tissue structure was initially simulated by generating a clustered point pattern using the Matern Cluster Process (Matérn 2013) with the following parameters:

- The intensity of the clusters: *κ* = 40
- The average number of cells per cluster: *µ* = 50
- The radius of each cluster: *r* = 0.1

Next, to capture the structure and density of the point pattern a kernel smoothed intensity (Diggle 1985) was calculated using sigma = 0.05. The density was thresholded by removing the bottom 25% of values to create a mask of the simulated tissue structure. Points were simulated as a Poisson point process in the regions inside and outside the mask to create the cell type C and cell type B respectively. To create cell type A, which exists on the boundary of cell type B and C, the product of the densities of B and C were taken and normalised creating a density which exists at the overlap between cell type B and C. A Poisson point process model was used on this density to create cell type A. In addition to this image, a context independent and random relationship image were simulated. In the context independent image, cell type A was creating using the density of cell type C. To create the random relationship image, all cell types were simulated using a Poisson point process model with an intensity of 12000 points per unit area. To each image a Kontextual curve was constructed.

### Survival analysis with Kontextual

To construct Kaplan Meier curves on the selected Keratin+tumour CD8 relationship, the Kontextual and L-function values were split by the median value, and the curves were plotted using the ggsurvfit R package (Sjoberg et al. 2023). Kontextual and L-function values were calculating using *r* = 100. The reported p-values were obtained from a Cox proportional hazard regression model on the original feature vector. To construct the bubble plot, Cox proportional hazard regression models were fit on all each pairwise Kontextual relationship (*r* = 100). The coefficients of each model were plotted on a bubble plot using the bubblePlot function.

When performing survival prediction, images containing “cold” tumours (n=5) were removed, due to sparse immune cell populations, creating unstable spatial features. Patient 22 and 38 were also filtered out due to missing survival information. In addition to the biologically informed Kontextual features we also used Kontextual with parent populations obtained from treekoR (Chan et al. 2021), the L-function, cell type proportions and cell type proportions relative to parent obtained using treekoR. All Kontextual and L-function values were calculated using *r* = 100. Kontextual and L-function values containing NAs or −100 were replaced with zeros, as they represented scenarios where the cell types being evaluated were not in the data or the populations were sparse and there were no cell types within the radius.

To perform survival prediction a regularised Cox Regression model was used, performance was assessed using 10 fold 100 repeat cross validation where the top 20 features determined by a Cox proportional hazard regression model were selected every fold. To evaluate performance Harrell’s C-index was used, which measures the concordance between predicted survival outcomes and the true survival outcomes. Classification and cross validation was performed using the ClassifyR package (Strbenac et al. 2015).

