## Supplementary figures for "Kontextual: Reframing analysis of spatial omics data reveals consistent cell relationships across images"

| Methods | Reference | Type | Observed value | Permuted mean | Permuted SD | Z-score |
|---|---|---|---|---|---|---|
| Kontextual | This study | Radii | 11.09 | 0.45 | 0.71 | 14.95 |
| L-function | (Besag 1977) | | -29.44 | 0.26 | 0.70 | -42.67 |
| Inhomogeneous L-function | (Baddeley et al. 2000) | | -21.93 | 0.04 | 0.65 | -21.99 |
| SpicyR | (Canete et al. 2022) | | -26.24 | 0.04 | 0.44 | -60.19 |
| kNN | (Windhager et al. 2023) | Graph | 1.05 | 2.50 | 0.03 | -51.90 |
| Expansion | | | 0.02 | 0.13 | 0.00 | -15.74 |
| Delaunay | | | 0.02 | 0.13 | 0.00 | -15.83 |
| DIMPLE | (Masotti et al. 2023) | Density | -0.33 | -0.05 | 0.03 | -7.91 |

**Table S1:** Several spatial analysis methods applied on the patient 6 image to assess the relationship between $P53^+$ tumour cells and immune cells. To assess the deviation from complete spatial randomness the mean and standard deviation were calculated by permuting cell type labels 1000 times. The Z-score was calculated by subtracting the permuted mean from the observed value and dividing by the standard deviation. A negative (positive) Z-score indicates dispersion (localisation) between the cell types.

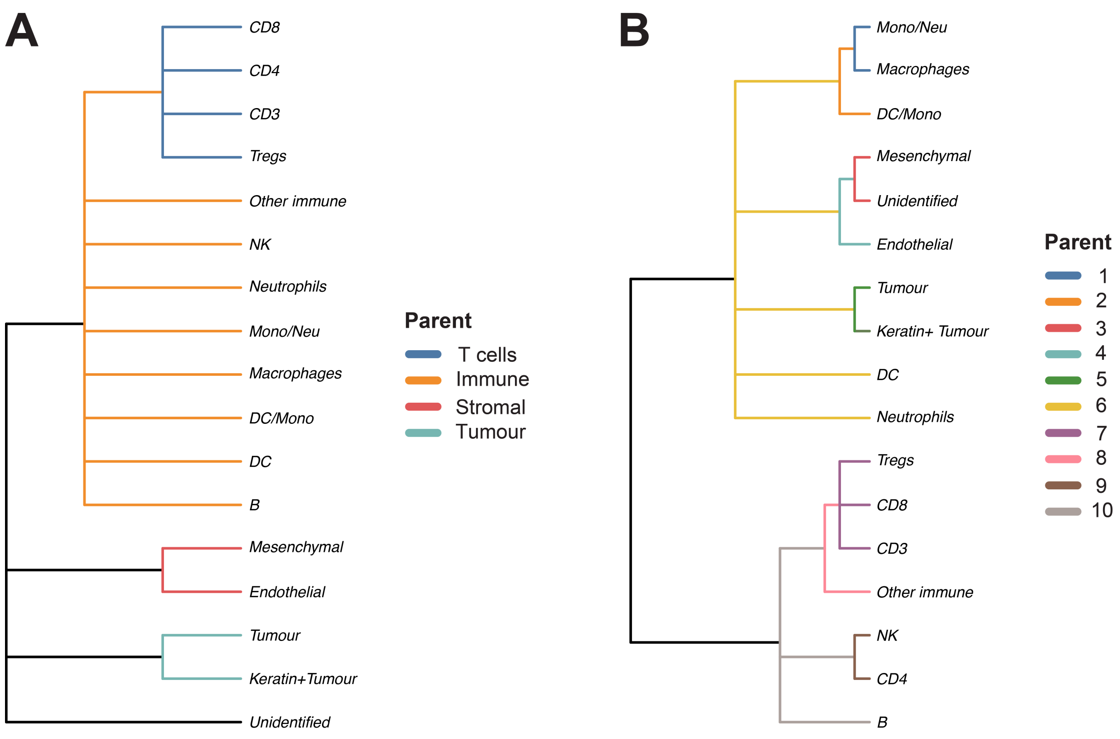


**Supplementary Figure S1:** Hierarchical cell structures used by Kontextual for Keren et al.'s breast cancer data. (**a**) Cell type hierarchy constructed using our biological understanding of the cell types, where cell types with shared generalised functions are grouped together. (**b**) Cell type hierarchy constructed with a HOPACH clustering strategy using the treekoR R package (Chan et al. 2021)

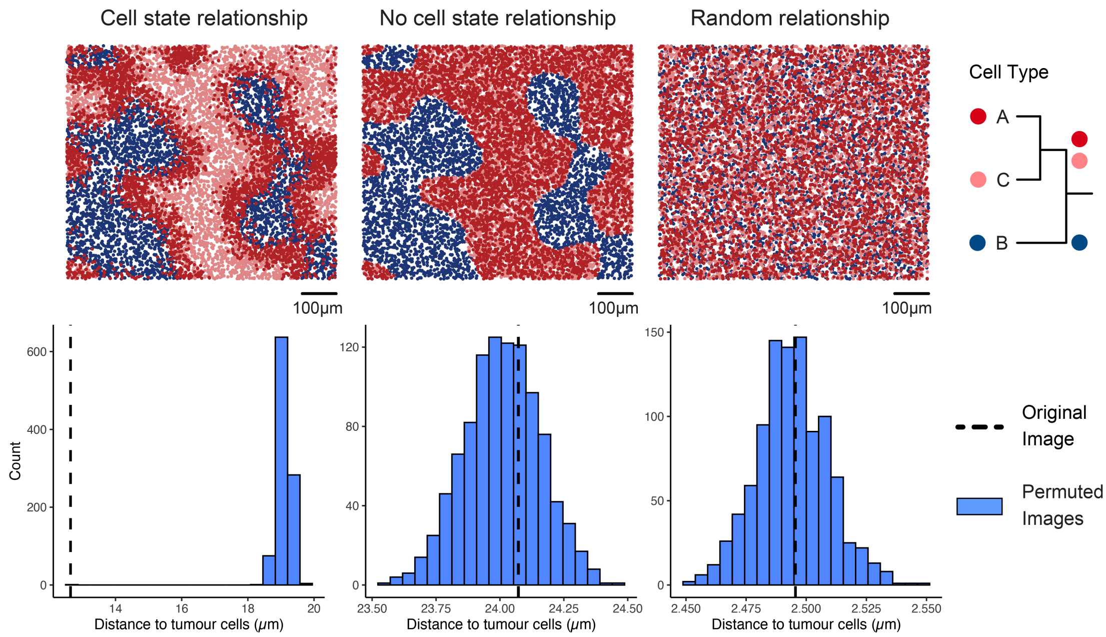


**Supplementary Figure S2:** Comparing the observed average distance of cell type A to their nearest cell type B (black line) compared to all of parent A (blue histogram) in the simulated images.
